# The sialome of the retina, alteration in age-related macular degeneration (AMD) pathology and potential impacts on Complement Factor H

**DOI:** 10.1101/2025.03.09.642149

**Authors:** Jaclyn Swan, Christopher B. Toomey, Max Bergstrand, Hector A Cuello, Jesse Robie, Hai Yu, Yue Yuan, Anoopjit Singh Kooner, Xi Chen, Jutamas Shaughnessy, Sanjay Ram, Ajit Varki, Pascal Gagneux

## Abstract

**Purpose:** Little is known about sialic acids of the human retina, despite their integral role in self/non-self-discrimination by complement factor H (CFH), the alternative complement pathway inhibitor.

**Methods:** A custom sialoglycan microarray was used to characterize the sialic acid-binding specificity of native CFH or recombinant molecules where IgG Fc was fused to CFH domains 16-20 (contains a sialic acid-binding site), domains 6-7 (contains a glycosaminoglycan-binding site) or the CFH-related proteins (CFHRs) 1 and 3. We analyzed macular and peripheral retinal tissue from post-mortem ocular globes for amount, type, and presentation (glycosidic linkage type) of sialic acid in individuals with age-related macular degeneration (AMD) and age-matched controls using fluorescent lectins and antibodies to detect sialic acid and endogenous CFH. Released sialic acids from neural retina, retinal pigmented epithelium (RPE) cells and the Bruch’s membrane (BrM) were labelled with 1,2-diamino-4,5-methylenedioxybenzene-2HCl (DMB), separated and quantified by high-performance liquid chromatography (DMB-HPLC).

**Results:** Both native CFH and the recombinant CFH domains 16-20 recognized Neu5Ac and Neu5Gc that is α2-3-linked to the underlying galactose. 4-*O*-Actylation of sialic acid and sulfation of GlcNAc did not inhibit binding. Different linkage types of sialic acid were localized at different layers of the retina. The greatest density of α2-3-sialic acid, which is the preferred ligand of CFH, did not colocalize with endogenous CFH. The level of sialic acids at the BrM/choroid interface of macula and peripheral retina of individuals with AMD were significantly reduced.

**Conclusions:** The sialome of the human retina is altered in AMD. This can affect CFH binding and consequently, alternative complement pathway regulation.

## Introduction

Limited information exists regarding the complex carbohydrates (glycans) present in the human retina, despite their crucial role in immune system regulation and physiology.^1^ Age-related macular degeneration (AMD) is the leading cause of blindness in individuals over the age of 60.^2^ Dysregulation of the alternative complement system contributes to the pathology of this disease.^3, 4^ The glycocalyx that coats the outside of cells, provides one of the first points of contact for cell-to-cell communication and defines molecular patterns on each cell. As these glycans are involved in self- and non-self-recognition, alteration of glycosylation has the potential to modulate interactions with immune regulatory proteins such as complement factor H (CFH), which is known to interact with negatively charged cell surface glycans including sialic acids and select glycosaminoglycans^5^, and malondialdehyde epitopes that result from oxidative stress.^6^ Previous studies have explored changes in IgG glycosylation and serum N-glycans as potential biomarkers for AMD^7^ but glycosylation changes within the retina have not been investigated. Understanding the normal and abnormal glycosylation patterns of the human retina with a focus on sialic acid will provide much needed insight into this multi-factorial disease.

Glycans are found conjugated to proteins, lipids and even RNA.^8^ Apart from providing a source of energy, this class of biomolecules also provides structural integrity, influences confirmation and activity of proteins and contributes to defining the molecular identity of cells and tissues. Glycans are attached to biomolecules through the enzymatically controlled glycosylation processes. In humans and other vertebrates most glycans carry terminal sialic acid. The molecular pattern formed by sialic acids defines the sialome a key component of the molecular frontier of each cell. Unlike proteins in which the sequence and structure can be predicted based on the genome, glycosylation is not template driven. Instead, glycosylation is the result of the combined action of several gene products, such as glycosyltransferases and glycosidases, and the availability of their substrates, activated monosaccharides (nucleotide sugars). For this reason, small environmental and biological changes can have dramatic effects on glycosylation. To add further complexity, glycans can be branched and different monosaccharides can be joined by different glycosidic linkage types in different positions, giving rise to immense molecular diversity, much of it with distinct biological characteristics. Glycans play essential roles in the cellular organization and development of multicellular organisms. For example, loss of sialic acid from the glycans of serum glycoproteins and red blood cells functions as an indicator of aging and signals for their removals from the circulatory system. Similarly, it is hypothesized that overall tissue sialyation also changes with age.

Despite some glycan analyses of the cornea and tear fluid, less is known about the glycan composition of the retina. The corneal epithelium contains densely O-glycosylated glycoproteins, specifically mucin (MUC16).^9^ The O-glycans on this mucin are heavily sialylated and hydrated, providing essential boundary lubrication and protection against potential pathogens. We essentially all see through a complex layer of glycoproteins. Alterations in the O-glycans of keratinized cells have been linked to inflammatory conditions such as superior limbic keratoconjunctivitis. Moreover, increased sulfation on O-glycans has been observed in the tear fluid of patients with ocular rosacea.^10, 11^ Existing retinal glycans studies have utilized lectin staining methods, employing glycan-binding proteins (lectins) to reveal the presence of sialic acids on drusen, indicated by positive staining with the lectin *Limax flavus* agglutinin (LFA).^12^ Drusen are the characteristic lesions observed in AMD, located between the Bruch’s membrane (BrM) and retinal pigmented epithelium (RPE) cells. Drusen exhibit minimal staining for terminal galactose (revealed by peanut agglutinin (PNA)). Treatment with sialidase to remove sialic acid results in decreased LFA staining and increased staining with PNA, suggesting that drusen are rich in sialic acid, predominantly linked to β1-3 linked galactose.^12, 13^ Sialic acid can be linked to underlying galactose or *N*-acetylgalactosamine in α2-3 and/or α2-6 linkage, or attached to another sialic acid with an α2-8 linkage, forming di-, oligo, or polysialic acids. LFA is believed to bind sialic acid regardless of its linkage to the underlying monosaccharide. Sialic acid is a key cell surface molecule in vertebrates where it provides most of the negative charge on cell surfaces. It is very important for biomedicine as humans have evolved distinct sialic acid biology and sialic acids form one of the primary motifs used by our innate immune system to distinguish self from non-self/altered-self.^14^

Multiple immune-regulating lectins within the human innate immune system discern self from non-self by detecting sialic acid. In some cases, the specific linkage of sialic acid is crucial for this recognition process. For example, CFH recognises α2-3 linked sialic acid.^15, 16^ CFH is a large, secreted glycoprotein and a key regulator of the alternative complement cascade. Dysregulation of the complement system is believed to be an underlying mechanism associated with AMD and the Y402H polymorphism in CFH accounts for one of the strongest risk factors demonstrating a potential role in the pathology.^17, 18^ CFH is made up of 20 short census repeat (SCR) domains, each containing approximately 60 amino acids joined by short linkers. CFH discriminates self from non-self via two separate glycan-binding domains. SCR6-8 is responsible for glycosaminoglycan (GAG) recognition^19^, whereas the SCR20 recognizes sialic acid with the specific linkage of α2-3, as well as GAGs.^5, 16^ In addition to recognising ‘self’ glycan motifs, CFH simultaneously inhibits the alternative pathway by accelerating the decay of C3 convertase (C3bBb) or acting as a co-factor for factor I cleavage of C3b to its hemolytically inactive form, iC3b. Recently, two inhibitors of the complement cascade have been shown to delay the progression of geographical atrophy in patients with advanced AMD.^20–22^ However, the efficacy and side-effect profile limits there use in clinical practice. Alternatively, alteration of the sialome (the complete repertoire of sialic acid types and linkages from a particular cell, tissue, or organism) has the potential to suppress overactive immune responses, including complement activation. Thus, we sought to determine whether there is a change in sialylation of human retina tissue in individuals with AMD.

Little is known about the retinal glycans in the normal aged eye and even less in degenerative conditions like AMD. Due to the non-regenerative nature of retinal cells and the plasticity of glycosylation, changes in glycan composition are possible correlates or even causes for immune dysregulation. Presentation of sialic acid in certain linkages on the termini of glycolipids, N- and O-glycans contributes to the cells, and in combination, the tissues presentation of ‘self’. Changes in presentation of sialic acid (linkage) or/and change in the quantity of sialic acid could both have the potential to contribute to AMD pathology. We aimed to further elucidate CFH specificities in recognising sialic acid using a sialoglycan array. Additionally post-mortem human ocular samples were used to investigate the distribution of sialic acid linkage through the neural retina and sub retina and the amount of sialic acid in the neural retina, RPE and BrM/choroid interface was quantified.

## Methods

### Sialoglycan array printing

The sialoglycan arrays were printed as described previously.^23^ Briefly, the source plate was prepared by aliquoting 20 µl of 10 mM chemically synthesised sialoglycans (each containing a terminal amino group on the aglycon) diluted in printing buffer (300 mM phosphate buffer pH 8.4) per well according to the plate template. The sialoglycan arrays were printed using ArrayIt SpotBot extreme instrument at 20°C 65% humidity, onto PolyAn 3D-NHS (Automate Scientific, PO-10400401).

Once slides were dried, they were blocked in pre warmed tris ethanolamine buffer (0.1 M Tris and 0.05 M ethanolamine, pH 9, 50°C) for 60 min gently rocking. Subsequently, immerse in pre-warmed Milli-Q water (50°C), followed by gentle rocking for 10 minutes in water at the same temperature. The water rinse and wash were then repeated a second time. The slides were dried by centrifuging in a horizontal slide holder for 10 min at 700 rpm. The slides were vacuumed sealed in a non-transparent container and stored at 4°C until used.

### Ocular dissection

Human ocular globes were collected from the San Diego Eye Bank with written consent from individual or next of kin. Due to the individuals being unidentifiable the collection of this tissue was under IRB exception from UC San Diego Human Research Protection Program. The human tissue experiments complied with the guidelines of the ARVO Best Practices for Using Human Eye Tissue in Research (Nov2021). Ocular samples were collected from males and females aged between 55-99 years of age (Supplementary Table S1). The anterior segment of the globe was first removed and samples, including 2 mm punches beside the fovea within the macular region and a 2 mm punches of the peripheral retina, were taken and snap frozen in optimal cutting temperature compound (OCT, Tissue Tek, Sakura Finetek, Torrance, CA, USA) to be used for cryosectioning. These were used for immunohistochemistry. Encompassing the 2mm punch in each area an 8 mm punch was taken. From this 8 mm punch the neural retina was removed and the presence of AMD pathology recorded. The RPE cells were carefully brushed off, collected, washed and snap frozen. Following this the BrM and choroid were peeled off from the sclera and as much of the choroid as possible was removed from the BrM.

### Histology

Cryosections from the stored OCT blocks were cut at 8 µm thick using a Leica CM1860 at −18°C and placed on to Superfrost™ Plus microscope slides (Fisherbrand™, 12-550-15). Each slide had six sections, with three sections from the macular and three sections from the peripheral retina. All slides were stored at −80°C. Prior to use the slides were defrosted at room temperature and air dried for a minimum of 1 hr and each frozen tissue section was circled using a PAP pen, to be able to concentrate the reagent correctly and prevent desiccation. Three individuals with AMD and five age matched controls were included in the histology experiments (Table 1). A haematoxylin and eosin (H&E) stain was standard in order to confirm tissue morphology. A summary of each of the assays performed on the slides with the corresponding controls is included in Table 2.

**Table 1.** An overview of the sialoglycan array assays conducted.

| Lectin | Primary antibody | Secondary antibody |
| --- | --- | --- |
| Native CFH (Quidel A410) (10 µg/ml) | Goat poly clonal anti-human CFH (0.5 µg/ml) | Anti-goat Cy5 (5 µg/ml) |
| rCFh16-20/mouseIgG2aFc (0.5 µg/ml) | Goat poly clonal anti-human CFH (0.5 µg/ml) | Anti-goat Cy5 (5 µg/ml) |
| rCFH6-10/mouseIgG2aFc (10-0.5 µg/ml) | Anti-mouse IgG Cy3 (5 µg/ml) | N/A |
| rCFHR1/humanIgG1Fc (5-0.5 µg/ml) | Anti-human IgG Cy3 (5 µg/ml) | N/A |
| rCFHR3/humanIgG1Fc (5-0.5 µg/ml) | Anti-human IgG Cy3 (5 µg/ml) | N/A |

**Table 2.** The lectins used to probe the histology slides and the corresponding controls and secondaries used.

| Primary protein | Source | Enzyme control | Secondary |
| --- | --- | --- | --- |
| SNA-FITC | VectorLabs | Sialidase | N/A |
| SGRP<sub>3</sub><sup>Hsa</sup> | <sup>23</sup> | Sialidase and<br>SGRP<sub>3</sub><sup>Hsa</sup>NB | Streptavidin Cy3 |
| mAb735 | <sup>24</sup> | EndoN | Anti-mouse IgG Cy3 |
| Goat polyclonal Anti-human CFH | <sup>25</sup> | N/A | Anti-goat IgG Cy3 |

### Immunofluorescence using SGRP3^Hsa^ for α2-3-linked sialic acid and SNA for α2-6-linked sialic acid

*Arthrobacter ureafaciens* sialidase (AUS; Roche, 10269611001) was diluted to 5 mU/ml in 50 mM sodium acetate buffer pH 5.2. Mock treatment included the sodium acetate buffer alone. The mock and sialidase solution were each overlaid on the necessary slides and incubated in a humid chamber overnight at 37°C. The slides were washed three times in PBS. All slides were fixed in 10% buffered formalin for 10 min and then wash. Autofluorescence was quenched using 1x TrueBlack (Biotum, 23007) solution in 70% ethanol (v/v), washed and avidin biotin blocked using the Vector labs kit (SP-2001). Prior to the addition of the primary lectin, all slides were overlayed with diluting buffer. Sialoglycan Recognizing Probe 3 (SGRP3*^Hsa^*) which is a sialic acid probe specific for α2-3-linked sialic acid was originally derived from *Streptococcus gordonii* and SGRP3*^Hsa^*non-binding (SGRP3*^Hsa^*NB) (carbohydrate binding mutant)^23^ were then added at 20 µg/ml but the directly conjugated SNA-FITC (Vector Labs NC9785207) was added at 5 µg/ml in diluting buffer. The slides were incubated for 1 hr at room temperature, washed and binding detected using streptavidin-Cy3 (Jackson ImmunoResearch, 109-160-084). All slides were washed and overlayed with nuclear Hoechst solution and then mounted with aqueous mounting media for viewing and photomicrography.

### Immunofluorescence using Anti-PolySialic acid

After fixation in 10% buffered formalin, slides were washed. Relevant slides were treated with enzyme activated recombinant EndoN enzyme 7 µg/ml in 2% BSA (w/v) for two hrs at 37°C or blocked in 2% BSA (w/v). Endogenous fluorescence was removed and mAb735 was diluted 1:1000 in 2% BSA and incubated for two hrs at room temp in a humid chamber. Slides were then washed and overlayed with anti-mouse IgG-Cy3 (Jackson ImmunoResearch, 115-165-071) for 45 min and then washed. All slides were treated with nuclear Hoechst stain and mounted using aqueous mounting media for viewing and digital photomicrography.

### Immunofluorescence using Anti-CFH

For the detection of endogenous CFH, the cryosections were initially fixed in ice cold acetone for 20 seconds and then washed in PBS. Autofluorescence was quenched using 1x TrueBlack in 70% ethanol for 1 min and then washing the slides in PBS. The slides were overlayed with diluting buffer (0.5% cold water fish gelatin in PBS) followed by goat polyclonal anti-human CFH diluted 1:500 in diluting buffer, incubated for 2 hrs at room temperature. After washing, binding was detected using Anti-goat Cy3 (Cy™3 AffiniPure Donkey Anti-Goat IgG (H+L) (min X Ck, GP, Sy Hms, Hrs, Hu, Ms, Rb, Rat Sr) Product Code: 705-165-147) diluted 1:500 in diluting buffer for 30 min. The nuclei were counter stained with Hoechst and then slides mounted with aqueous mounting media for viewing and digital photomicrography.

### Imaging

All microscopy imaging was captured on a Keyence Biorevo BZ-9000 microscope. Post-processing such as brightening and merging was conducted with Fiji 2.14.0/1.54F.

### Sialic acid identification and quantification (DMB-HPLC)

Ocular tissue was suspended in Milli-Q water with the addition of 1x Halt protease inhibitor cocktail (Thermo Scientific, 1861278) and 1mM PMSF. The tissue was homogenized by sonication in 15-second bursts at approximately 15% intensity, followed by a 45-second rest, repeated three times or until the tissue was fully homogenised. Protein concentration was determined using a BCA assay (Thermo Scientific, 23225). Tissue lysate was stored at −20°C until use the following day.

### Acid release of sialic acids

The starting material type dictated the quantity of protein from which the sialic acid was released. Approximately, 10 µg of BrM lysate, 20 µg of neural retina lysate and as much RPE cell lysate as possible was incubated in 2 M acetic acid at 80°C for 3 hrs, gently shaking at 300 rpm. The released sialic acid was separated from the lysate by passing through a 10 kDa spin filter. The flow through was frozen at −80°C and lyophilised. The free sialic acid was resuspended in 100 µl of Milli-Q water. An aliquot of this was base treated to remove the O-acetyl groups by adding NaOH to a final concentration of 0.1 M and incubating at 37°C for 1 hr. The pH was subsequently adjusted back to approximately pH 7 by dropwise addition of HCl. The resuspended acid released sialic acid and the base treated sialic acid were fluorescently labelled by the addition of 7 mM 1,2-diamino-4,5-methylenedioxybenzene dihydrochloride, 1.6 M glacial acetic acid, 0.75 M β-mercaptoethanol and 18 mM sodium hydrosulphite at a 1:1 ratio. The labelling occurred in the dark at 4°C for 48 hrs to prevent loss and migration of *O*-acetyl modifications as described previously.^26^ Standards including, 1 nmol of Neu5Ac and free sialic acid from bovine submaxillary mucins (BSM) was also labelled in parallel.

Depending on the concentration of sialic acid in the sample, either the maximum amount of labelled sialic acid or 50% was injected into the LaChrome Elite Hitachi HPLC system with a Phenomenex Gemini C18 column (5 μm, 250 mm × 4.6 mm) at room temperature. A gradient was run over 68 mins switching from solution A (acetonitrile: methanol: water, 7:7:86 (v/v/v)) to solution B (acetonitrile: methanol: water, 11:7:82 (v/v/v)) with an additional 7 min of 100% solution B with a consistent flow rate of 0.9 ml/min.^26^ The DMB-labelled sialic acid species were detected a fluorescence detector at an excitation of 373 nm and emission of 448 nm.

Using the Neu5Ac standard to generate a standard curve, the amount of Neu5Ac in each sample was determined, and the concentrations of Neu5Ac were normalized to the protein concentration of the original sample. The types of sialic acid present were determined by comparison to the BSM standard. In Prism graph pad multiple unpaired t-test with Welch correction and no assumption of variance of each group were performed.

## Results

### Sialic acid-binding specificity of complement factor H (CFH)

It was evident that rCFH16-20 is very specific for sialic acid in an α2-3-linkage to galactose (Fig. 1A). Similar to previous results, it binds both Neu5Ac and Neu5Gc^16, 27^ but did not recognize other sialic acid species such as 2-keto-3-deoxy-D-glycero-D-galactonononic acid (Kdn) or legionaminic acid (Leg). However, there was variability in binding depending on the linkage and monosaccharide attached to the reducing terminal of Siaα3Gal and included N-/O-glycan sialoglycan motifs and ganglioside type motifs. For examples, Neu5Ac/Neu5Gcα3Galβ4GlcNAc and their various modifications (9OAc, 4OAc, 4NAc and 6S) are commonly identified on N- and O-glycans. Recombinant CFH was also found to strongly interact with Neu5Gcα3Galβ3GalNAc, which is representative of core 1 O-glycans, and its counterparts containing a 9OAc or 4OAc-modified Neu5Gc. CFH also recognized, GM3 ganglioside-type glycans, Neu5Ac/Neu5Gcα3Galβ4Glc and modifications with 4OAc and 4NAc at the sialic acid. However, recognition was lost if the sialic acid was capped with an α2-8-linked sialic acid (GD3). The addition of β1-4 GalNAc to create GM2 also prevented the recognition of CFH. Similarly, the presence of fucose in Lewis X structures [Neu5Acα3Galβ4(Fucα3)GlcNAc] prevented the recognition of CFH. Interestingly, CFH did not recognize any 7,9-diOAc sialic acid species. In all cases just the sialylgalactose disaccharide alone was enough to initiate CFH binding. This suggests that CFH does not specifically recognize one class of glycans.

**Figure 1.**
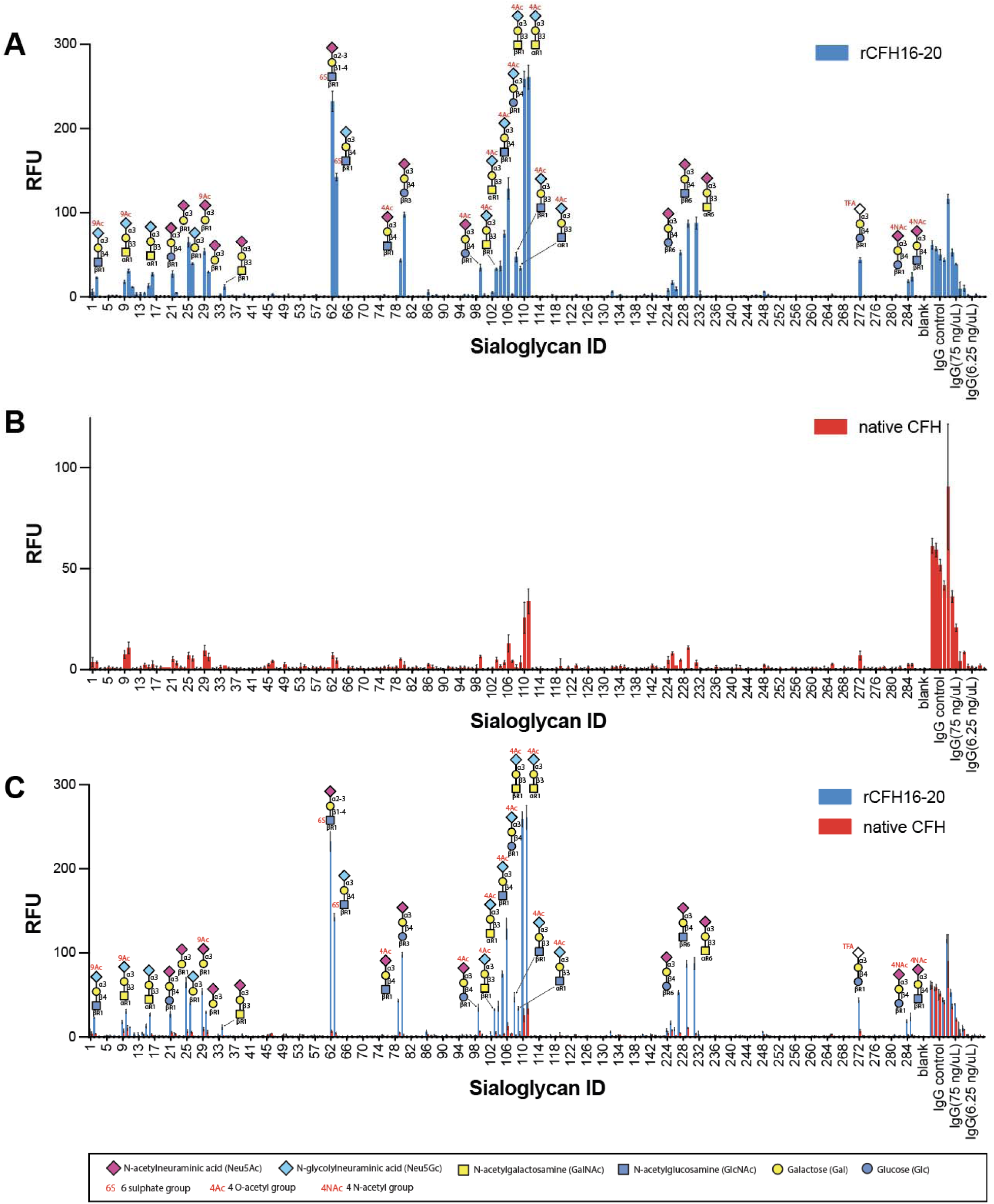
Human complement factor H (CFH) recognition of sialic acid. (A) Recombinant CFH domains 16-20 fused to Fc (rCFH16-20) (0.5 µg/ml) the composition of the sialogycans recognized are depicted. (B) native CFH purified fro human serum (10 µg/ml). (C) an overlay to compare the recognition of rCFH16-20 and native CFH. Relative fluorescent units (RFU). Error bars represent standard deviation of four replicate spots. For a list of all glycan compositions included on the array see Supplementary Table S2.

A sulphated GlcNAc (sialoglycan #62) showed a greater amount of rCFH16-20 binding than its non-sulphated counterparts. Furthermore, rCFH16-20 showed a preference for 4-*O*-acetylated Neu5Gc although binding to 4-*O*-acetylated Neu5Ac was also observed.

The recombinant CFH16-20 and native full length CFH, purified from human serum, bound the same sialoglycans (Fig. 1B), although the dominant interaction for native CFH was with Neu4,5Ac_2_3Galβ3GalNAc (sialoglycan #110 and #111). The affinity of the full-length protein appeared to be much lower than that of the truncated recombinant protein on the glycan array (Fig. 1C). The recombinant CFH16-20 was used at 0.5 µg/ml and the native CFH as used at 10 µg/ml. The difference in the number of sialic acid binding site does not account for the difference in the amount of binding. The dimeric nature of the rCFH16-20 molecule, on account of it being expressed as an Fc fusion protein, may have contributed to its greater affinity compared to the monomeric full-length CFH.

Recombinant GAG binding domain of CFH, rCFH6-10, was passed over the sialoglycan micro array at several concentrations however no binding was observed. Similarly, recombinant CFHR1 or CFHR3 fused to Fc did not bind to any of the sialoglycans on the microarray (data not shown).

### Histology

#### H&E

Due to the nature of the collection and the fragility of the retina tissue, mechanical damage artifacts and retinal detachment was commonly observed during collection and cryosectioning. In some cases, the outer and inner photoreceptor segments were missing and in other cases the RPE was dislocated as was evident in the H& E staining (Fig. 2A).

**Figure 2.**
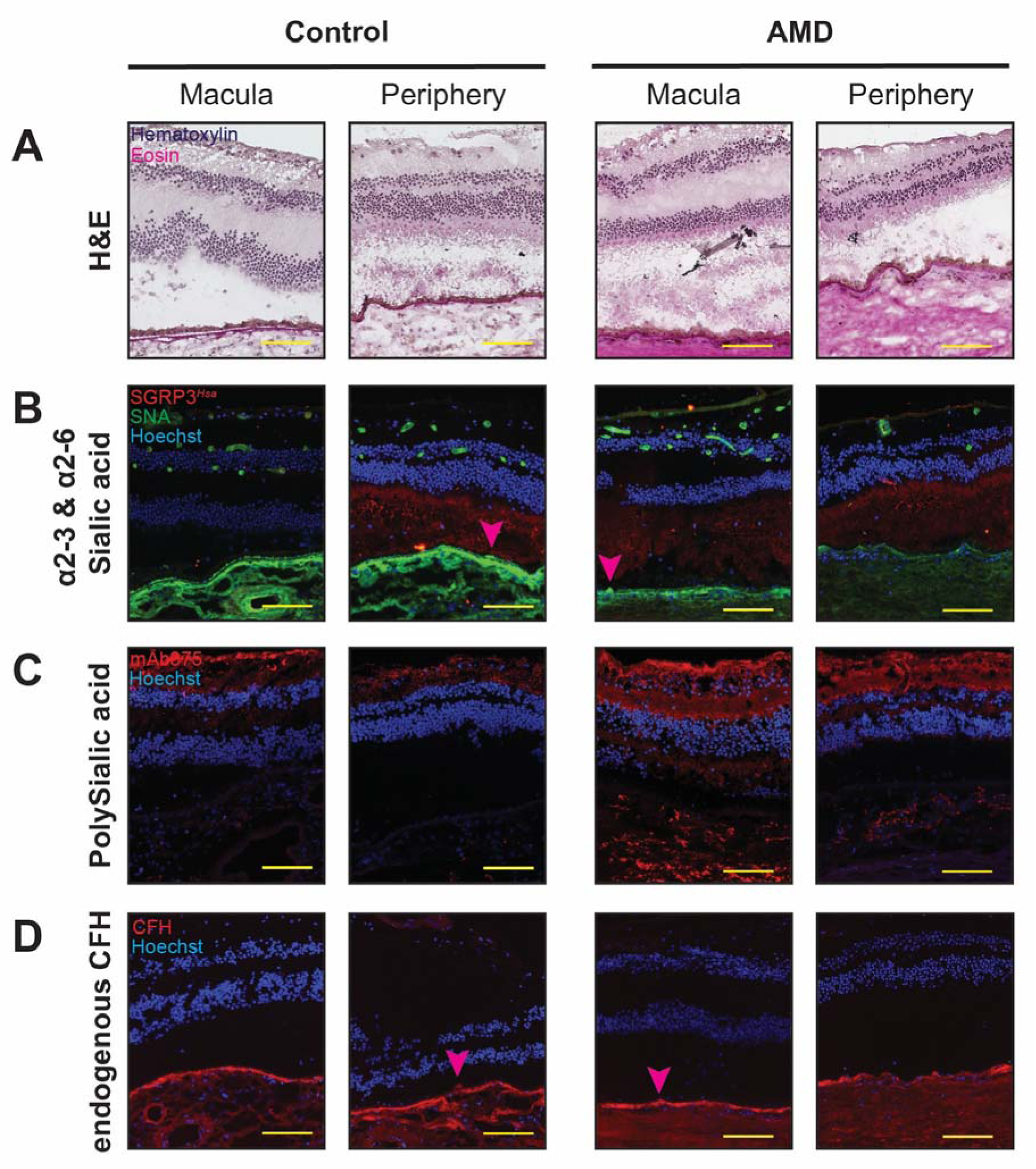
The distribution of different sialic acid linkages throughout the retina of individuals with and without age related macular degeneration (AMD). (A) Haematoxylin and eosin stain (H&E). (B) SGRP3*^Hsa^* recognition of α2-3-linked sialic acid and SNA α2-6-linked sialic acid. (C) Monoclonal antibody 735 (mAb735) binds α2-8 polysialic acid. (D) Endogenous CFH detected with goat polyclonal anti-human CFH detecting endogenous CFH. The yellow scale bar is 100µm. Pink arrowhead indicated drusen. Comparison of individual variation can be observed in Supplementary Fig. S1.

#### Detection of α2-6 and α2-3 sialic acid in human retinas

SNA is a lectin produced in the bark of *Sambucus nigra* (European elder berry tree), it recognises a variety of sialic acid species but strictly linked to the underlying glycan in an α2-6-sialyl linkage. Commonly SNA is used to stain the epithelium of blood vessels throughout the body, the use of SNA in this study clearly illustrated the vasculature in the inner layers of the retina, the choriocapillaris and choroidal vasculature (Fig. 2B). Moderate detection of α2-6-linked sialic acid was observed in the BrM and choroid. Interestingly, individual number 3 showed strong detection of α2-6-linked sialic acid on the inner as well as outer side of the BrM (Supplementary Figure S1). Strong staining of the drusen in two individual was observed (Fig. 2B, pink arrowhead). Very little detection of α2-6-linked sialic acid was observed on the RPE cells of all individuals. No difference was observed in detection of α2-6-linked sialic acid between macula and peripheral regions of AMD individuals or controls. Sialidase treatment appropriately abolished binding of SNA, however in the mock-treated samples a comparable amount of binding was observed (Supplementary Figure S2), demonstrating that none of the SNA binding was due to non-specific protein-protein interactions. The sialic acid probe specific for α2-3-linked sialic acid (SGRP3*^Hsa^*) was originally derived from *Streptococcus gordonii.* SGRP3*^Hsa^* bound to the photoreceptors very specifically (Fig 2B). The amount of binding to the photoreceptors in macula and peripheral region was variable for each individual, however there was no specific trend. Unfortunately, after overnight sialidase and mock treatment, the majority of the SGRP3*^Hsa^* binding was lost but the SNA binding remained unchanged (Supplementary Figure S2). Potentially the long exposure to the low pH and temperature could have hydrolysed some of the sialic acid. However, no recognition was observed for the non-binding SGRP3*^Hsa^* which has a mutated sialic acid binding domain. TrueBlack successfully quenched the autofluorescence observed due to the accumulation of lipofuscins in the RPE cells as no signal was detected in any of the negative controls.

#### Detection of Poly sialic acid in human retinas

Polysialic acid was identified and mapped in the human retina. This was achieved using a monoclonal mouse antibody (mAB735) developed against colominic acid (α2-8 polysialic acid) produced by a strain of pathogenic *E. coli* K1 bacterium. These results were corroborated by staining with SGRP8 which also recognizes α2-8-linked sialic acid (SGRP8 data not shown). As a negative control, the tissue section was treated with an active recombinant EndoN enzyme, which specifically cleaves α2-8-linked sialic acid, to demonstrate the mAb was specific for polysialic acid identified in the human retina (Supplementary Figure S3). The distribution of α2-8 linked sialic acid appears to be very specific, for the innermost layers of the retina (ganglion cell layers and inner plexiform layer) (Fig. 2C). However, in two of the three AMD individuals evaluated strong detection of α2-8-linked sialic acid was observed in localized area of the choroid (Fig. 2C).

#### Detection of Endogenous CFH in human retinas

The presence of endogenous CFH was observed throughout the choroid and sclera for all individuals in tissue collected from both the macula and peripheral regions (Fig. 2D). Very strong detection of endogenous CFH was seen in the choriocapillaris of most individuals. Although detection in the choroidal vasculature was most prominent in individual number 1 (Supplementary Figure S1). Consistent with previous work^28^, endogenous CFH was detected on drusen. Considerable inter-individual variation was observed and there was no difference between AMD and controls, and no obvious trend when comparing peripheral tissue to macula. No endogenous CFH was detected in the neural retina of any of the individual regardless of AMD status or region the tissue was taken from, suggesting systemic CFH cannot cross the blood retina barrier and if RPE or neural retina cells produce CFH, it is below the level of detection.

### Comparison of sialic acid content in the sialome of AMD patients (DMB-HPLC)

The amount of sialic acid in the neural retina, RPE cells and the BrM/choroid interface was measured by releasing sialic acid with mild acid, labeling with DMB followed by HPLC separation. Each sample was analyzed in comparison to bovine BSM standard (dark blue chromatogram; Fig. 3A) and base treated (red chromatogram; Fig. 3A) to determine if any *O*-acetylated sialic acid species were present. A very small amount of 7-*O*-acetylated and 9-*O*-acetylated Neu5Ac was detected in some of the samples. However, this proportion made up less than 1% of the total sialic acid. As expected, no Neu5Gc was detected (humans have lost the capacity to produce this sialic acid derived from Neu5Ac due to a loss-of function mutation of the gene encoding the enzyme CMAH, which is now fixed in the human population).

**Figure 3.**
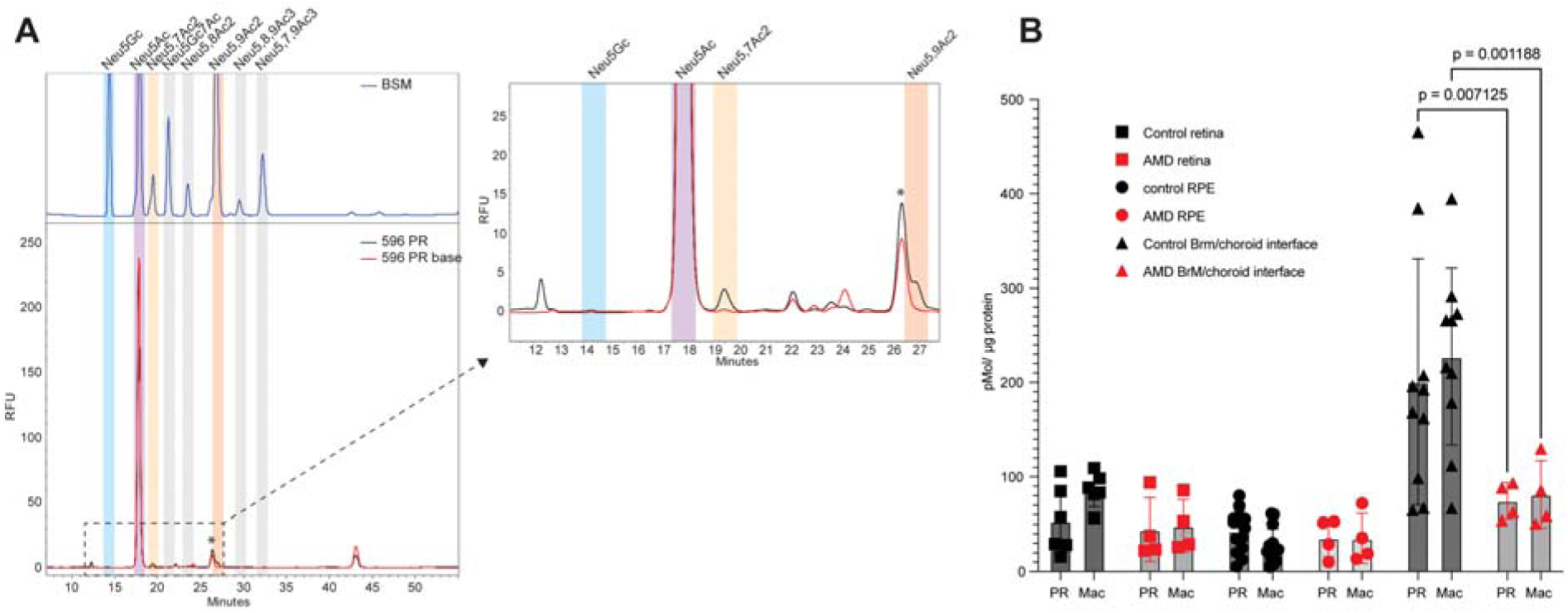
Identification and quantification of sialic acid in the human retina with and without age related macular degeneration (AMD). (A) Representative DMB-HPLC profile. Retention time (RT) of peaks detected in the retinal samples (lower panel, black) are compared to RT of well-characterized distinct types of sialic acids present in bovine submaxillary mucins (BSM) (upper panel, blue). Additionally, the released sialic acids are base treated (lower panel, red) to remove any potential *O*-acetyl groups, confining the modification of the sialic acid. (B) The quantity of Neu5Ac in each sample from both the peripheral retina (PR) and macula (Mac) was determined by comparisons to standards of known concentration. An unpaired t-test with Welch correction and no assumption of variance of each group, error bars represent the standard error of the mean.

For each batch, a standard curve was generated by running known concentration of Neu5Ac standards, enabling the quantification of Neu5Ac in each sample. Subsequently, the Neu5Ac levels were normalized to the initial protein concentration. There was no difference between the amount of Neu5Ac in the macula compared to the peripheral retina for any of the different retinal tissues collected. However, a lower amount of sialic acid was detected in the neural retina and the RPE cells compared to the BrM/choroid interface (Fig. 3B). Although there was individual variation, there is no significant difference between the amount of sialic acid between individuals with and without AMD in the neural retina and RPE cells. However, individuals with AMD had significantly less sialic acid in the BrM/choroid interface compared to aged-matched individuals in both the macula and peripheral retina (p= 0.02 and p=0.004, respectively) (Fig. 3B).

## Discussion

To our knowledge this is the first indepth study of the human ocular sialome in combination with the distribution of endogenous CFH protein. Utilizing a sialoglycan array has illustrated the intricacies of the siaclic acid-binding domain of CFH recognition for different presentations of sialic acid, with various modifications. Eyes from healthy subjects and AMD patients revealed variation in sialic acid linkage across the different layers of the retina. Lastly, the quantity of sialic acid in BrM/choroid interface is significantly reduced in individuals who had AMD, in both the macula and peripheral region of the retina. This suggests that sialic acid may play a role in the organization of the retina and in AMD pathology.

Through the use of a diverse sialoglycan array it was confirmed both native CFH and rCFH16-20 specifically bind α2-3-linked sialic acids, which is consistent with previous literature.^15, 16^ However, a preference for 4-*O*-acetylated sialic acid was observed, which has not been demonstrated before. Each of the sialoglycans that had previously shown weak but positive interaction with full-length CFH (in the micromolar and millimolar range) through the saturation transfer difference (STD) nuclear magnetic resonance (NMR) studies was identified on the sialoglycan array.^15, 16^ Unexpectedly, interactions with 9-*O*-acetylated Neu5Ac and 9-*O*-acetylated Neu5Gc were observed, contrary to previous findings which demonstrated increased CFH binding to murine erythroleukemia cells following treatment with a 9-*O*-acetylesterase^29^ and markedly decreased CFH binding to lacto-N-neotetraose (LNnT) in gonococcal lipooligosaccharide (LOS) capped with Neu5,9Ac_2_ compared to LOS bearing Neu5Ac on LNnT.^27^ However, this difference could be due to the way the modified sialic acid is presented on biological surfaces, where it may interact with other surface molecules, versus in purified form on microarrays. Additionally, the presence of additional CFH ligand such as C3b/C3d^30^ or PorB.1A^31^ (both non-glycan mediated) have previously been demonstrated to enhance avidity, this level of complexity is lacking from this assay. As demonstrated in the past, it does not appear that human CFH has any particular preference over Neu5Ac or Neu5Gc.^16^ Similarly, the sialic acid binding domain of many Siglecs has rapidly evolved allowing Siglecs to preference both Neu5Ac and Neu5Gc, instead of only recognizing Neu5Gc which is the case for non-human primate Siglecs ^32, 33^. Humans are unable to synthesize Neu5Gc from its precursor Neu5Ac and the source of Neu5Gc on any human tissue is due to dietary uptake and metabolic incorporation.^34^ Therefore, the CFH recognition of Neu5Gc could reflect an ancestral condition in human evolution, from when our ancestors still possessed a functional CMP-Neu5Ac hydroxylase gene, required for the synthesis of Neu5Gc.^35, 36^

The glycan array also demonstrated that sulfation of the GlcNAc in the LacNAc motif underlying the terminal sialic acid increased recognition of CFH. Similar fine-tuning of sialic acid binding has also been observed for Siglec 3 (CD33, sialic acid-binding Ig superfamily lectin 3) another immune modulating sialic acid receptor.^37^ This is the first time that CFH has been shown to recognize 4-*O*-acetylated sialic acid. Previously, Neu4,5Ac_2_ has been found in horses, guinea pigs, monotremes and several fish species but its presence in humans has not been definitively established to date.^38–41^ Although, Neu4Ac5Gc has been detected in human patients with colon cancer.^42^ Therefore, it is interesting that CFH, a self-recognition complement regulator is recognizing this modified version of sialic acid. The distribution of this sialic acid modification is likely to be underrepresented in the literature due to the difficulty of its detection by HPLC and lack of commercially available probes or antibodies.^43^ It will be interesting to speculate what role this interaction could possibly be having considering its distributions in humans is predicted to be minimal, if not absent.

It is important to consider that CFH does not recognize sialic acid in isolation, the glycocalyx and extra cellular matrix consists of many different types of glycans and glycoconjugates. Their density and specific combinations could each play a role in CFH recognition (even other glycans which do not directly interact^44^). Additionally, CFH can recognize both sialic acid and GAGs, the relationship between sialic acid and GAG recognition of CFH has not been investigated. Technically CFH could recognize both sialic acid and GAGs at the same time through simultaneous interactions with both SCR 7 and SCR 18-20, and the effect this dual interaction may have on protein function or surface avidity is unknown. It is conceivable that the sialome could change but then be compensated by the GAGome and therefore have little effect on CFH recognition.

The retina is a highly organised tissue, consisting of distinct layers, each defined by different cell types. We believe the different cell types in each layer is what is contributing to the very specific localisation of each of the different sialic acid linkage types throughout the tissue. The only region where there appears to be considerable overlap is in the BrM and choriocapillaris that appear to be highly sialylated with both α2-3 and α2-6-linkages, but no polysialic acid. Although not apparent in the histology, the amount of sialic acid present in the neural retina and RPE cells is significantly less than what was detected in the BrM/choroid interface, demonstrating the advantage of combining histology with biochemical analysis of the analytes. Interestingly none of the sialic acid linkage-specific lectins recognized Muller cells, which span the whole neural retina. The photoreceptors appear to have a very distinct sialome consisting of only α2-3-linked sialic acid. Previous lectin studies have also demonstrated that the photoreceptors and more specifically the cone photoreceptors stain very strongly with PNA suggesting an abundance of uncapped O-glycans.^45^

CFH has been shown to bind to oxidized lipids and bisretinoids on drusen.^46^ Although intense staining of SNA and, to some extent, SGRP3*^Hsa^* was observed on the drusen, along with previous LFA staining^12^, the interaction between CFH and drusen is not necessarily a glycan-protein interaction. Oxidative stress leads to the formation of malondialdehyde, increasing the interaction of CFH and preventing pro-inflammatory events.^6, 46, 47^ This interaction occurs at the polyanion binding site of CFH within SCR 19-20, which could compete with negatively charged glycans for binding.^47^ We found that the localization of endogenous CFH does not align with the areas showing the highest staining for α2-3-linked sialic acid in the photoreceptors. However, our results also illustrate that systemic CFH does not appear to be crossing the blood retina barrier, although accumulation on and in drusen was observed. The amount of CFH reported to be produced by the RPE cells must be below the level of detection by our specific antibodies or in line with current presumptions CFH produced by RPE is secreted basally. Despite the change in sialic acid observed in the BrM, a change in the amount of endogenous CFH was not observed. However, the goat polyclonal antibody is not capable of differentiating between full length CFH and the truncated splice variant CFH like (FHL) which consists only of SCR 1-7. It also cross reacts with CFHR1 and CFHR3 (CFHRs (2, 4 and 5) have not been tested). FHL and the CFHRs can compete with CFH for binding to surfaces. Therefore, variable expression of each of these CFH family members could affect complement regulation as only the full length CFH and FHL are capable of inhibiting the alternative complement system.^48^

Although the amount and type of sialic acid in the neural retina and RPE cells does not change between individuals with AMD and those without, there is a significant difference in quantity of sialic acid in the BrM/choroid interface. Lack of terminal sialic acid is thought to be a marker for complement activation and apoptosis.^49^ Interestingly, in individuals without AMD the quantity of sialic acid in the BrM/choroid interface is much greater than other regions of the retina. In individuals with AMD this reverts to a level comparable with the neural retina and RPE cells. Whether there is increased sialidase activity in this region or downregulation of sialyltransferases is yet to be determined. In a previous model of uveitis, another inflammatory eye disease, an upregulation of NEU1 in Muller cells has been reported.^50^ The factors contributing to the reduced quantity of Neu5Ac in the BrM is currently unknown. Possible explanations include a reduction in sialylated N-glycans, O-glycans or gangliosides. Many proteins which make up the BrM are potentially glycosylated including collagen, laminin and integrins.^51–53^ Unfortunately, this reduction in sialic acid quantity in the BrM/choroid interface was not observable in the histology. How this change in the sialome could affect AMD pathology remains speculative. To date, the role of sialic acid in the blood-retina/brain-barrier remains unclear. It would be interesting to know if sialome alteration could induce leakage of this crucial selectively permeable layer. However, cell bound, rapidly evolving sialic acid-binding lectins, the Siglecs, which are expressed in different combinations on most immune cells, are likely to be involved in differential responses to altered sialomes.^54^ We hypothesize that a reduced amount of sialic acid would result in less inhibition of microglia (Siglec 1, 3, 11 and 16) and macrophages (Siglec 3, 7 and 9), however immune regulation was beyond the scope of this study.^54, 55^ Recently, Gne+/− knockout mice, with reduced amounts of the key enzyme in sialic acid synthesis, showed reduced protein-bound polysialic acid in the neural retina.^56^ This led to increased lysophagosomal activity in retinal microglia, upregulation of pro-inflammatory IL-1β, and loss of rod bipolar cells. Knocking out complement factor C3 partially reversed these effects, underscoring the connection between sialic acid and the alternative complement system.^56^ Nanoparticles decorated in polysialic acid are currently in human trials.^57^ It is believed to inhibit microglia/macrophage by interacting with Siglecs.^58^

The GAG-binding domain of CFH did not interact with any sialoglycans on the array even those with an α2-8-linkage. A recent study showed a weak interaction between polysialic nanoparticles and CFH and CFH domains 6-7 fused to Fc.^59^ The polysialic acid nanoparticles could enhance CFH binding to C3b in surface plasmon resonance assays and modestly reduced hemolytic activity of the alternative and classical pathways of complement.^59^ This contrasts with our data and prior studies.^59^ While this study and prior studies have not shown a direct interaction between CFH and α2-8 polysialic acid or the ability of colominic acid to enhance the interaction between CFH and C3b^60, 61^, differences in experimental outcomes may have resulted from differences in the formulations of colominic acid (i.e., nanoparticles versus soluble colominic acid), the concentration of ligands used (experiments with polysialic acid nanoparticles used ligand concentrations two to three orders of magnitude higher than prior studies) and/or the type of assay used. Nevertheless, the reduction in the quantity of sialic acid in the BrM of individuals with AMD seen in this study warrants further study of sialic acid-based treatment options that compensate for the reduction of sialic acid in disease eyes.

This study provides novel insight into the spatial distribution of sialylated glycans in normal and AMD-affected eyes. Using multiple lectins and controlling endogenous auto fluorescence has illustrated clear localisation of each of the different sialic acid linkages. A decrease of sialic acid in the BrM/choroid interface of individuals with AMD suggested a change in sialome could be contributing to the progressive disease. Although only a small number of individuals were included in this study due to the limitation of deriving post-mortem human ocular tissue.

## Commercial Relationships Disclosure

N/A

## Acknowledgements

We would like to thank Professor Nissi Varki for all her advice regarding the histology component of this project and many helpful discussions.

## References

1. Varki A. Biological roles of glycans. Glycobiology 2016;27:3–49.

2. Mitchell P, Liew G, Gopinath B, Wong TY. Age-related macular degeneration. Lancet 2018;392:1147–1159.

3. Wilke GA, Apte RS. Complement regulation in the eye: implications for age-related macular degeneration. J Clin Invest 2024;134.

4. de Jong S, Tang J, Clark SJ. Age-related macular degeneration: A disease of extracellular complement amplification. Immunol Rev 2023;313:279–297.

5. Ram S, Sharma AK, Simpson SD, et al. A novel sialic acid binding site on factor H mediates serum resistance of sialylated Neisseria gonorrhoeae. J Exp Med 1998;187:743–752.

6. Weismann D, Hartvigsen K, Lauer N, et al. Complement factor H binds malondialdehyde epitopes and protects from oxidative stress. Nature 2011;478:76–81.

7. Bućan I, Škunca Herman J, Jerončić Tomić I, et al. N-Glycosylation Patterns across the Age-Related Macular Degeneration Spectrum. Molecules 2022;27:1774.

8. Flynn RA, Pedram K, Malaker SA, et al. Small RNAs are modified with N-glycans and displayed on the surface of living cells. Cell 2021;184:3109–3124.e3122.

9. Brockhausen I, Elimova E, Woodward AM, Argüeso P. Glycosylation pathways of human corneal and conjunctival epithelial cell mucins. Carbohydr Res 2018;470:50–56.

10. Ozcan S, An HJ, Vieira AC, et al. Characterization of novel O-glycans isolated from tear and saliva of ocular rosacea patients. J Proteome Res 2013;12:1090–1100.

11. Watanabe H, Maeda N, Kiritoshi A, Hamano T, Shimomura Y, Tano Y. Expression of a mucin-like glycoprotein produced by ocular surface epithelium in normal and keratinized cells. Am J Ophthalmol 1997;124:751–757.

12. Mullins RF, Johnson LV, Anderson DH, Hageman GS. Characterization of drusen-associated glycoconjugates. Ophthalmology 1997;104:288–294.

13. Mullins RF, Hageman GS. Human ocular drusen possess novel core domains with a distinct carbohydrate composition. J Histochem Cytochem 1999;47:1533–1539.

14. Varki A. Letter to the glyco-forum: since there are PAMPs and DAMPs, there must be SAMPs? Glycan “self-associated molecular patterns” dampen innate immunity, but pathogens can mimic them. Glycobiology 2011;21:1121–1124.

15. Blaum BS, Hannan JP, Herbert AP, Kavanagh D, Uhrín D, Stehle T. Structural basis for sialic acid–mediated self-recognition by complement factor H. Nat Chem Biol 2015;11:77–82.

16. Schmidt CQ, Hipgrave Ederveen AL, Harder MJ, Wuhrer M, Stehle T, Blaum BS. Biophysical analysis of sialic acid recognition by the complement regulator Factor H. Glycobiology 2018;28:765–773.

17. Klein RJ, Zeiss C, Chew EY, et al. Complement factor H polymorphism in age-related macular degeneration. Science 2005;308:385–389.

18. Toomey CB, Johnson LV, Bowes Rickman C. Complement factor H in AMD: Bridging genetic associations and pathobiology. Prog Retin Eye Res 2018;62:38–57.

19. Prosser BE, Johnson S, Roversi P, et al. Structural basis for complement factor H linked age-related macular degeneration. J Exp Med 2007;204:2277–2283.

20. Liao DS, Grossi FV, El Mehdi D, et al. Complement C3 inhibitor pegcetacoplan for geographic atrophy secondary to age-related macular degeneration: a randomized phase 2 trial. Ophthalmology 2020;127:186–195.

21. Heier JS, Lad EM, Holz FG, et al. Pegcetacoplan for the treatment of geographic atrophy secondary to age-related macular degeneration (OAKS and DERBY): two multicentre, randomised, double-masked, sham-controlled, phase 3 trials. Lancet 2023;402:1434–1448.

22. Khanani AM, Patel SS, Staurenghi G, et al. Efficacy and safety of avacincaptad pegol in patients with geographic atrophy (GATHER2): 12-month results from a randomised, double-masked, phase 3 trial. Lancet 2023;402:1449–1458.

23. Srivastava S, Verhagen A, Sasmal A, et al. Development and applications of sialoglycan-recognizing probes (SGRPs) with defined specificities: exploring the dynamic mammalian sialoglycome. Glycobiology 2022;32:1116–1136.

24. Frosch M, Görgen I, Boulnois GJ, Timmis KN, Bitter-Suermann D. NZB mouse system for production of monoclonal antibodies to weak bacterial antigens: isolation of an IgG antibody to the polysaccharide capsules of Escherichia coli K1 and group B meningococci. Proc Natl Acad Sci U S A 1985;82:1194–1198.

25. Agarwal S, Ram S, Ngampasutadol J, Gulati S, Zipfel PF, Rice PA. Factor H facilitates adherence of *Neisseria gonorrhoeae* to complement receptor 3 on eukaryotic cells. J Immunol 2010;185:4344–4353.

26. Ji Y, Sasmal A, Li W, et al. Reversible O-Acetyl Migration within the Sialic Acid Side Chain and Its Influence on Protein Recognition. ACS Chem Biol 2021;16:1951–1960.

27. Gulati S, Schoenhofen IC, Whitfield DM, et al. Utilizing CMP-Sialic Acid Analogs to Unravel Neisseria gonorrhoeae Lipooligosaccharide-Mediated Complement Resistance and Design Novel Therapeutics. PLoS Pathog 2015;11:e1005290.

28. Bhutto IA, Baba T, Merges C, Juriasinghani V, McLeod DS, Lutty GA. C-reactive protein and complement factor H in aged human eyes and eyes with age-related macular degeneration. Br J Ophthalmol 2011;95:1323–1330.

29. Shi W-X, Chammas R, Varki NM, Powell L, Varki A. Sialic acid 9-O-acetylation on murine erythroleukemia cells affects complement activation, binding to I-type lectins, and tissue homing. J Biol Chem 1996;271:31526–31532.

30. Kajander T, Lehtinen MJ, Hyvärinen S, et al. Dual interaction of factor H with C3d and glycosaminoglycans in host–nonhost discrimination by complement. Proc Natl Acad Sci U S A 2011;108:2897–2902.

31. Ngampasutadol J, Ram S, Gulati S, et al. Human factor H interacts selectively with Neisseria gonorrhoeae and results in species-specific complement evasion. J Immunol 2008;180:3426–3435.

32. Sonnenburg JL, Altheide TK, Varki A. A uniquely human consequence of domain-specific functional adaptation in a sialic acid–binding receptor. Glycobiology 2003;14:339–346.

33. Saha S, Khan N, Comi T, et al. Evolution of Human-Specific Alleles Protecting Cognitive Function of Grandmothers. Mol Biol Evol 2022;39.

34. Tangvoranuntakul P, Gagneux P, Diaz S, et al. Human uptake and incorporation of an immunogenic nonhuman dietary sialic acid. Proc Natl Acad Sci U S A 2003;100:12045–12050.

35. Chou H-H, Takematsu H, Diaz S, et al. A mutation in human CMP-sialic acid hydroxylase occurred after the *Homo-Pan* divergence. Proc Natl Acad Sci U S A 1998;95:11751–11756.

36. Irie A, Koyama S, Kozutsumi Y, Kawasaki T, Suzuki A. The Molecular Basis for the Absence of N-Glycolylneuraminic Acid in Humans*. J Biol Chem 1998;273:15866–15871.

37. Jung J, Enterina JR, Bui DT, et al. Carbohydrate sulfation as a mechanism for fine-tuning siglec ligands. ACS Chem Biol 2021;16:2673–2689.

38. Schauer R, Schmid H, Pommerencke J, Iwersen M, Kohla G. Metabolism and role of O-acetylated sialic acids. The Molecular Immunology of Complex Carbohydrates—2 2001;325–342.

39. Aamelfot M, Dale OB, Weli SC, Koppang EO, Falk K. The in situ distribution of glycoprotein-bound 4-O-Acetylated sialic acids in vertebrates. Glycoconj J 2014;31:327–335.

40. Kamerling JP, Dorland L, van Halbeek H, Vliegenthart JF, Messer M, Schauer R. Structural studies of 4-O-acetyl-alpha-N-acetylneuraminyl-(2 goes to 3)-lactose, the main oligosaccharide in echidna milk. Carbohydr Res 1982;100:331–340.

41. Urashima T, Inamori H, Fukuda K, Saito T, Messer M, Oftedal OT. 4-O-Acetyl-sialic acid (Neu4,5Ac2) in acidic milk oligosaccharides of the platypus (*Ornithorhynchus anatinus*) and its evolutionary significance. Glycobiology 2015;25:683–697.

42. Miyoshi I, Higashi H, Hirabayashi Y, Kato S, Naiki M. Detection of 4-O-acetyl-N-glycolylneuraminyl lactosylceramide as one of tumor-associated antigens in human colon cancer tissues by specific antibody. Mol Immunol 1986;23:631–638.

43. Manzi AE, Dell A, Azadi P, Varki A. Studies of naturally occurring modifications of sialic acids by fast-atom bombardment-mass spectrometry. Analysis of positional isomers by periodate cleavage. J Biol Chem 1990;265:8094–8107.

44. Cohen M, Hurtado-Ziola N, Varki A. ABO blood group glycans modulate sialic acid recognition on erythrocytes. Blood 2009;114:3668–3676.

45. Blanks JC, Johnson LV. Specific binding of peanut lectin to a class of retinal photoreceptor cells. A species comparison. Invest Ophthalmol Vis Sci 1984;25:546–557.

46. Meri S, Haapasalo K. Function and dysfunction of complement factor H during formation of lipid-rich deposits. Front Immunol 2020;11:611830.

47. Hyvärinen S, Uchida K, Varjosalo M, Jokela R, Jokiranta TS. Recognition of malondialdehyde-modified proteins by the C terminus of complement factor H is mediated via the polyanion binding site and impaired by mutations found in atypical hemolytic uremic syndrome. J Biol Chem 2014;289:4295–4306.

48. Cipriani V, Tierney A, Griffiths JR, et al. Beyond factor H: The impact of genetic-risk variants for age-related macular degeneration on circulating factor-H-like 1 and factor-H-related protein concentrations. Am J Hum Genet 2021;108:1385–1400.

49. Meesmann HM, Fehr EM, Kierschke S, et al. Decrease of sialic acid residues as an eat-me signal on the surface of apoptotic lymphocytes. J Cell Sci 2010;123:3347–3356.

50. Lorenz L, Amann B, Hirmer S, Degroote RL, Hauck SM, Deeg CA. NEU1 is more abundant in uveitic retina with concomitant desialylation of retinal cells. Glycobiology 2021;31:873–883.

51. Tvaroška I. Glycosylation Modulates the Structure and Functions of Collagen: A Review. Molecules 2024;29:1417.

52. Domogatskaya A, Rodin S, Tryggvason K. Functional diversity of laminins. Annu Rev Cell Dev Biol 2012;28:523–553.

53. Aisenbrey S, Zhang M, Bacher D, Yee J, Brunken WJ, Hunter DD. Retinal pigment epithelial cells synthesize laminins, including laminin 5, and adhere to them through alpha3- and alpha6-containing integrins. Invest Ophthalmol Vis Sci 2006;47:5537–5544.

54. Varki A, Angata T. Siglecs—the major subfamily of I-type lectins. Glycobiology 2005;16:1R–27R.

55. Linnartz-Gerlach B, Kopatz J, Neumann H. Siglec functions of microglia. Glycobiology 2014;24:794–799.

56. Cuevas-Rios G, Assale TA, Wissfeld J, et al. Decreased sialylation elicits complement-related microglia response and bipolar cell loss in the mouse retina. Glia 2024;72:2295–2312.

57. Krishnan A, Callanan DG, Sendra VG, et al. Comprehensive Ocular and Systemic Safety Evaluation of Polysialic Acid-Decorated Immune Modulating Therapeutic Nanoparticles (PolySia-NPs) to Support Entry into First-in-Human Clinical Trials. Pharmaceuticals 2024;17:481.

58. Krishnan A, Sendra VG, Patel D, et al. PolySialic acid-nanoparticles inhibit macrophage mediated inflammation through Siglec agonism: a potential treatment for age related macular degeneration. Front Immunol 2023;14:1237016.

59. Peterson SL, Krishnan A, Patel D, et al. PolySialic Acid Nanoparticles Actuate Complement-Factor-H-Mediated Inhibition of the Alternative Complement Pathway: A Safer Potential Therapy for Age-Related Macular Degeneration. Pharmaceuticals (Basel) 2024;17.

60. Meri S, Pangburn MK. Discrimination between activators and nonactivators of the alternative pathway of complement: regulation via a sialic acid/polyanion binding site on factor H. Proc Natl Acad Sci U S A 1990;87:3982–3986.

61. Meri S, Pangburn MK. Regulation of alternative pathway complement activation by glycosaminoglycans: specificity of the polyanion binding site on factor H. Biochem Biophys Res Commun 1994;198:52–59.

