## Supplementary figures and sup table 1 for "The sialome of the retina, alteration in age-related macular degeneration (AMD) pathology and potential impacts on Complement Factor H"

**Supplementary Material**

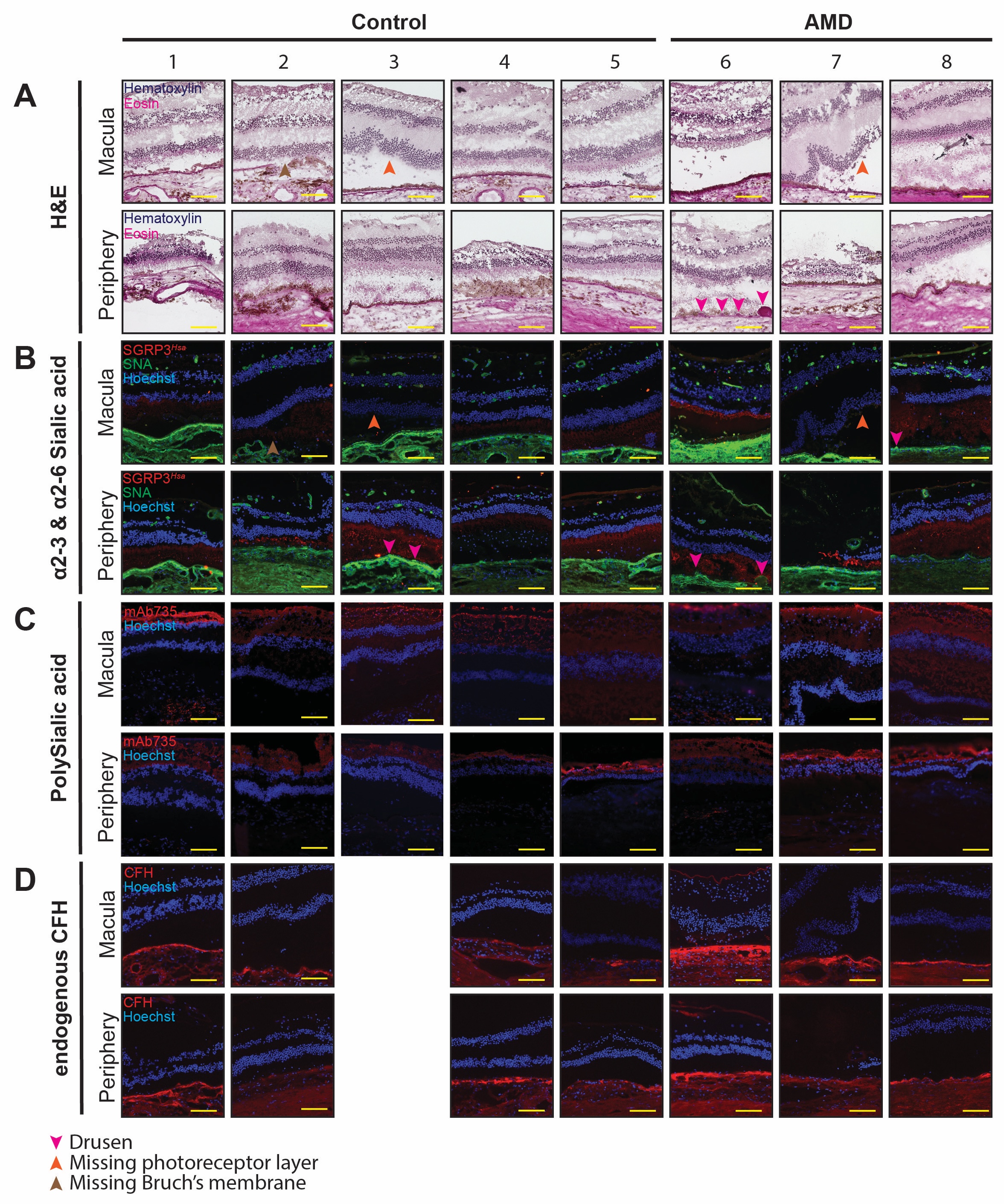

**Supplementary Fig. S1.** The distribution of different sialic acid linkages throughout the retina of individuals with (n=3) and without age related macular degeneration (AMD) (n=5). (A) Hematoxylin and eosin stain (H&E). (B) SGRP3*^Hsa^* recognizes α2-3-linked sialic acid and SNA α2-6-linked sialic acid. (C) Monoclonal antibody 735 (mAb735) binds α2-8 polysialic acid. (D) Endogenous CFH detected with goat polyclonal anti-human CFH detecting endogenous CFH. The yellow scale bar is 100µm. Pink arrowhead indicated drusen; orange arrowhead shows tissue with missing photoreceptors and the brown arrow heads indicates areas where the Bruch’s membrane is missing.

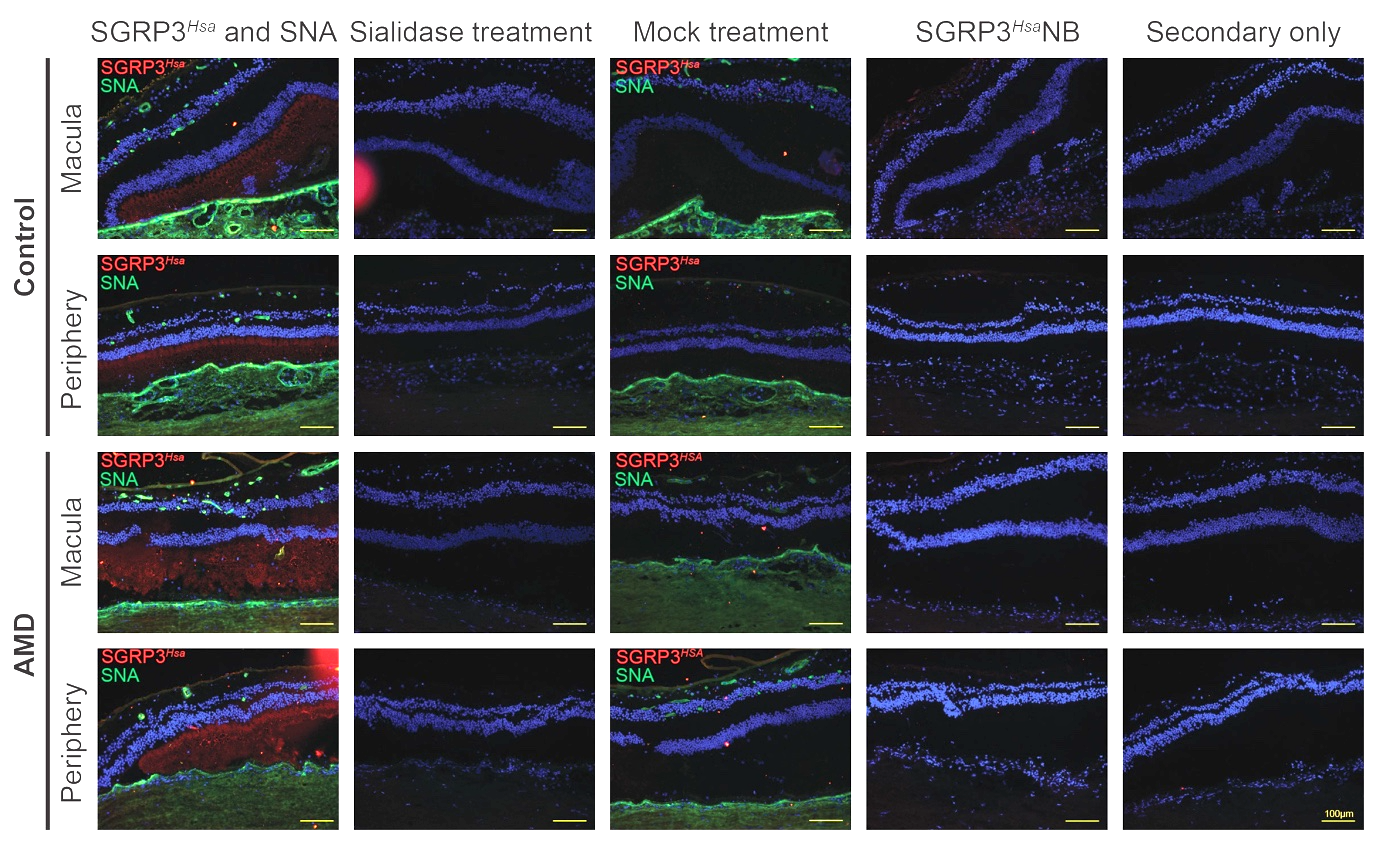
**Supplementary Fig. S2.** Multi-lectin staining and negative controls.

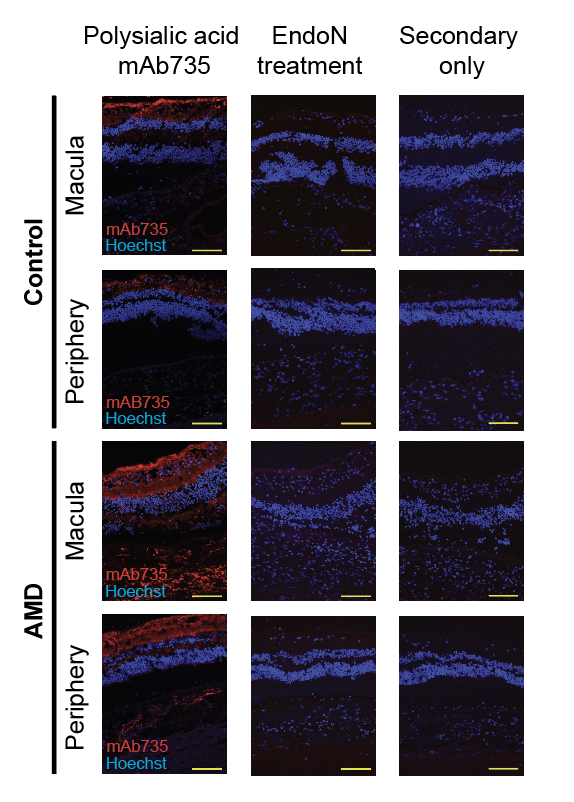

**Supplementary Fig.S3.** Polysialic acid staining using mAB735 and negative controls which include EndoN treatment to cleave polysialic acid and secondary only control. Yellow scale bar = 100 µm.

**Supplementary Table S1.** Details of the ocular donors and which experiments they were included in.

|  |  |  |  | **Death to refrigeration** | **Death to enucleation** | **Death to dissection** | **DMB-HPLC** | | |  | **Histology** | | | | **Cause of death** |
| --- | --- | --- | --- | --- | --- | --- | --- | --- | --- | --- | --- | --- | --- | --- | --- |
| **ID** | **sex** | **age** | **AMD status** | **(hrs:mins)** | | | **Retina** | **RPE** | **BrM/ choroid** | **H&E** | **SNA** | **SGRP3*^Hsa^*** | **mAb375** | **ahCFH** |  |
| 1 | M | 79 | no | 4:23 | 18:45 | 23:03 |  |  |  |  |  |  |  |  | Myocardial Infraction |
| 2 | M | 69 | no | 7:00 | 26:57 | 32:20 |  |  |  |  |  |  |  |  | Hypertensive Cardiovascular Disease |
| 3 | F | 93 | no | 2:45 | 5:45 | 8:57 |  |  |  |  |  |  |  |  | Dementia |
| 4 | F | 74 | no | 2:31 | 21:36 | 23:19 |  |  |  |  |  |  |  |  | Intraparenchymal hemorrhage |
| 5 | M | 80 | no | 6:45 | 17:23 | 19:15 |  |  |  |  |  |  |  |  | Myocardial Infraction |
| 6 | F | 69 | AMD | 10:22 | 11:58 | 15:22 |  |  |  |  |  |  |  |  | N/A |
| 7 | M | 79 | AMD | 1:53 | 23:23 | 28:22 |  |  |  |  |  |  |  |  | Pneumonia |
| 8 | M | 82 | AMD | 2:50 | 20:57 | 27:10 |  |  |  |  |  |  |  |  | Blunt Force Injury d/t Trauma -Peds vs Auto |
| 9 | F | 72 | no | 4:00 | 26:21 | 52:15 |  |  |  |  |  |  |  |  | Cerebral Vascular Accident |
| 10 | M | 73 | no | 4:00 | 23:08 | 47:25 |  |  |  |  |  |  |  |  | Cancer- Pancreas |
| 11 | f | 80 | no | n/a | 23:38 | 31:25 |  |  |  |  |  |  |  |  | Congrstive Heart Failure |
| 12 | M | 81 | no | 2:12 | 24:54 | 29:07 |  |  |  |  |  |  |  |  | Myocardial Infraction |
| 13 | F | 71 | no | 3:55 | 8:40 | 11:27 |  |  |  |  |  |  |  |  | Jaw Cancer |
| 14 | M | 73 | no | 3:59 | 8:29 | 11:52 |  |  |  |  |  |  |  |  | Sepsis |
| 15 | F | 69 | no | 8:29 | 13:56 | 16:20 |  |  |  |  |  |  |  |  | Cervical Cancer |
| 16 | M | 65 | no | n/a | 5:54 | 25:15 |  |  |  |  |  |  |  |  | Septic Shock |
| 17 | F | 99 | AMD | 2:39 | 13:23 | 16:16 |  |  |  |  |  |  |  |  | Sepsis |
